# Exploring distinct and joint contributions of the Locus Coeruleus and the Substantia Nigra/Ventral Tegmental Area complex to reward and valence processing using high-resolution fMRI

**DOI:** 10.1101/2024.01.31.577328

**Authors:** J. M. Hall, D. Shahnazian, R. M. Krebs

## Abstract

Dopaminergic neurons in the substantia nigra and the ventral tegmental area (SN/VTA) are classically viewed as key mediators in reward processing, while noradrenergic cells in the locus coeruleus (LC) are thought to modulate (negative) saliency processing. However, this conventional distinction is being revised by more recent research in animals. To explore the respective contributions of both the LC and SN/VTA in reward and valence processing in humans, we assessed fMRI data during stimulus encoding and response phase of a rewarded emotion-discrimination task (n=38). Participants responded significantly faster to reward-predicting and negative valence stimuli compared to their non-salient counterparts. LC activity was overall higher during trials involving reward prospect, and in particular for reward trials featuring positive valence, demonstrating an additive effect of reward and positive valence in LC. Moreover, LC activity was differentially increased for negative compared to positive valence in the response phase, indexing its role in invigorating responses to negative events. The SN/VTA showed increased activity in the response phase of reward trials (neutral valence) and negative valence trials (no-reward), which aligns with coding relative saliency of these events in their respective contexts. LC modulations were accompanied by covariations in occipital cortex, suggesting noradrenergic contributions to visual prioritization of salient events. The findings underscore the sensitivity of both LC and SN/VTA to reward prospects and negative valence, challenging the dominant view of SN/VTA’s involvement in merely positive events and emphasizing their essential role in action invigoration above and beyond mere stimulus encoding. The intricate roles of the DA and NA system in reward and emotional valence processing in humans warrant further exploration and validation, given the limitations inherent to neuroimaging of deep brain structures.

## 1. Introduction

Dopaminergic neurons of the substantia nigra pars compacta and the ventral tegmental area (referred to as SN/VTA) in the midbrain are classically considered key mediators of reward processing. Previous work has shown a consistent activation of the dopaminergic system in response to rewards and reward-predicting stimuli (Cohen et al., 2012; Schultz, 2002, 2013; Schultz et al., 1997; Steinberg et al., 2013). Moreover, animal as well as well as patient studies demonstrated that dopaminergic projections to the nucleus accumbens and frontal regions are vital for motivational processes (Aarts et al., 2011; Abler et al., 2006; K. C. Berridge & Robinson, 1998; Robbins & Everitt, 1992). Dopaminergic signalling in mesolimbic areas also increases during processing positive valence stimuli (for reviews, see Burgdorf & Panksepp, 2006; Wise, 2004). However, the function of dopamine (DA) may not be specific to processing rewarding or other positive events as increased DA activation is also observed following novel (no-reward) stimuli (Bunzeck & Düzel, 2006; Krebs et al., 2011), as well as negative, aversive 2003).

The source region of noradrenaline (NA), the locus coeruleus (LC) extensively projects to the neocortex and subcortical structures including the thalamus, septum, hippocampus and basal lateral amygdala (Loughlin et al., 1986). These widespread connections allow the NA system to regulate general arousal states by modulating neuronal gain (Aston-Jones & Cohen, 2005). More recently, it has become clear that the LC pathways are particularly selective (for review, see Sara & Bouret, 2012). For example, connectivity with the hippocampus and amygdala contribute to memory formation (McIntyre et al., 2012), whereas NA in the frontal areas is a key modulator of working memory, attention (Arnsten et al., 2012) and cognitive flexibility (Aston-Jones & Cohen, 2005; Bouret & Sara, 2005; Jahn et al., 2018).

Interestingly, previous work employing electroencephalography in humans suggested that the NA system is also sensitive to motivationally relevant stimuli (Nieuwenhuis et al., 2005). Consistent with these results, more recent animal studies have demonstrated that the NA system responds to reward-predicting cues (Bouret & Richmond, 2015; Jahn et al., 2018). For example, in an experimental task, four monkeys were presented with cues indicating whether a reward could be earned by timely releasing a bar (Bouret & Richmond, 2015). Both dopaminergic neurons in the SN and noradrenergic neurons in the LC showed transient activation at time of the cue, as well as during the release of the bar. Moreover, both SN and LC activity have been shown to increase when expecting a reward by coding value-based decision making (Varazzani et al., 2015). The magnitude of this increase in NA activity was also associated to effort, as measured by physical force produced. The authors suggested that the role of NA could be complementary to that of DA, in which NA by regulates effort to obtain the reward by mobilizing resources to energize behaviour. While DA may have a stronger influence on reward-based decisions, several studies have shown that DA can also promote approach actions in anticipation of an upcoming reward (Bouret et al., 2012).

More generally, it has been suggested that DA and NA may mediate a learning signal to optimize behaviour and both NA and DA activity may be modulated by overall saliency (Pessoa, 2009). Salient stimuli, whether they are affective, unexpected, or rewarding, strengthen representations and capture attention. While saliency might seem an obvious common denominator to trigger DA and NA responses, a complete functional overlap would seem highly inefficient and unlikely considering that dissociations have been reported for other more complex cognitive operations in humans that go beyond ad hoc stimulus processing.

The current study focused on the neural activity within the DA and NA during reward and valence processing in humans using high-resolution fMRI. To this end, we assessed activity in the SN/VTA and the LC regions while participants performed a rewarded emotion-discrimination paradigm. Importantly, in this task, both reward prospect (reward, no reward) and emotional valence (positive, neutral, negative) were similarly relevant, and factorially combined from trial to trial, which allowed to assess their potential interaction on the neural level. Further, stimulus presentation and response phase were separated by design to assess evaluative processes and action invigoration, respectively, mirroring related animal work (Bouret et al., 2012; Bouret & Richmond, 2015). We expected both the DA and NA systems to be activated in reward trials, and most prominently during response implementation that will eventually lead to the rewarding outcomes (Bouret et al., 2012). In addition, we expected modulations based on stimulus valence, with increased DA activation during positive stimuli (Bradley & Lang, 2007; Park et al., 2018), and a dissociation between the DA and NA system for negative valence, with a differential increase in NA activity (Berridge, 2008; McCall et al., 2017). No a priori hypotheses were formed about whether these valence modulations would be mostly expected during the stimulus presentation or the response phase. Finally, we explored potential interactions between reward and valence processing, which could take the form of additive effects (e.g., a positive stimulus might further increase the reward response) or in terms of relative salience (e.g., a negative stimulus might be more salient in a no-reward context).

## 2. Materials and methods

### 2.1.1 Participants

Based on our power analysis (see https://osf.io/2rsye), 44 participants were recruited through the Ghent University online recruiting website and social media websites. The inclusion criteria included (in addition to the safety screening for MRI participation) age between 18 – 35 years, no (history of) diagnosis of a neuropsychiatric disorder, (corrected) normal vision and right-handedness. Six participants were excluded from the data analyses (behavioural and fMRI) due to low task performance (predefined as <60% correct trials), data coding issues or motion artifacts that could not be corrected for using noise reduction techniques. The final sample consisted of 38 participants (mean age = 23.1 years; SD = 5.2; 22 (58%) females). Participants were reimbursed 40€ (plus an additional performance-dependent reward of up to 10€) for the entire session, which lasted approximately 90 minutes. The experimental procedures were approved by the Ghent University Hospital Ethics Committee, and all participants gave their written informed consent before the start of the study. This study was preregistered on the Open Science Framework (https://osf.io/2rsye).

### 2.1.2 Experimental paradigm

Participants performed a rewarded emotion-discrimination task in the fMRI scanner (Figure 1A). In each trial, a unique face stimulus (referred to as stimulus event) with negative (angry), positive (happy), or neutral emotional expression was presented for 1000 ms. The gender (male/female) of the stimulus signalled reward prospect (of 7 cents per correct answer), or no-reward prospect (reward/no-reward). The mapping between gender and reward prospect was counterbalanced across participants. Following this stimulus, two of the three response options (negative/positive/neutral) were shown in Dutch on the left and right side of the screen (referred to as response event). These corresponded to the three valence categories, i.e., angry (BOOS), neutral (NEUT), and happy (BLIJ). One word was matching the expression of the face stimulus, while one was not. Using an MR compatible response box, participants had to select the word that matched the emotional expression of the preceding face stimulus by pressing the left or right response button corresponding to the word location on screen with the index and middle finger of their right hand. Responses needed to be made within 1000 ms. The spatial location of the matching word was random to avoid any spatial biases for a specific valence category. Feedback was provided after each trial (referred to as feedback event) for 1000 ms indicating whether the answer was correct (check mark), incorrect (cross), or too late (clock symbol), and whether or not a reward had been obtained: after correct responses in reward trials, the bonus (“7 ct”) was displayed underneath the check mark. In all other cases (incorrect or too late reward trials, as well as all no reward trials), the feedback display included “0 ct”. All stimuli were presented on a black background. In between the stimulus event, the response event, and feedback event, as well as between trials, a white fixation cross was shown for a variable stimulus-onset-asynchrony (SOA) of 2090 – 8360 ms. These SOAs between the events were distributed based on multiples of the scan repetition time (TR), with 80% at 1 TR, 10% at 2 TRs, 5% at 3 TRs and 4 TRs, to allow for effective event-related BOLD response estimation across trials (Hinrichs et al., 2000). This experiment included stimuli from the NimStim face stimulus set (Tottenham et al., 2009) and included 72 happy, 72 neutral, and 72 angry facial expressions posed by 36 actors, who expressed all three emotions (and similar levels of arousal (Adolph & Alpers, 2010; Smith et al., 2013)). Participants completed three task blocks in the MR scanner (corresponding to three functional runs), which were programmed using PsychoPy version 2.3 (Python version 3.8) (Peirce, 2007). Each block consisted of 144 trials that lasted approximately 8 min. At the end of each block, the cumulative feedback about the total earned reward amount was shown. At the end of the experiment, each participant was paid their total amount of earned reward, in addition to the base reimbursement. Note that the project also included recognition memory data, which has not been considered here due to missing data and hence a lack of power.

**Figure 1.**
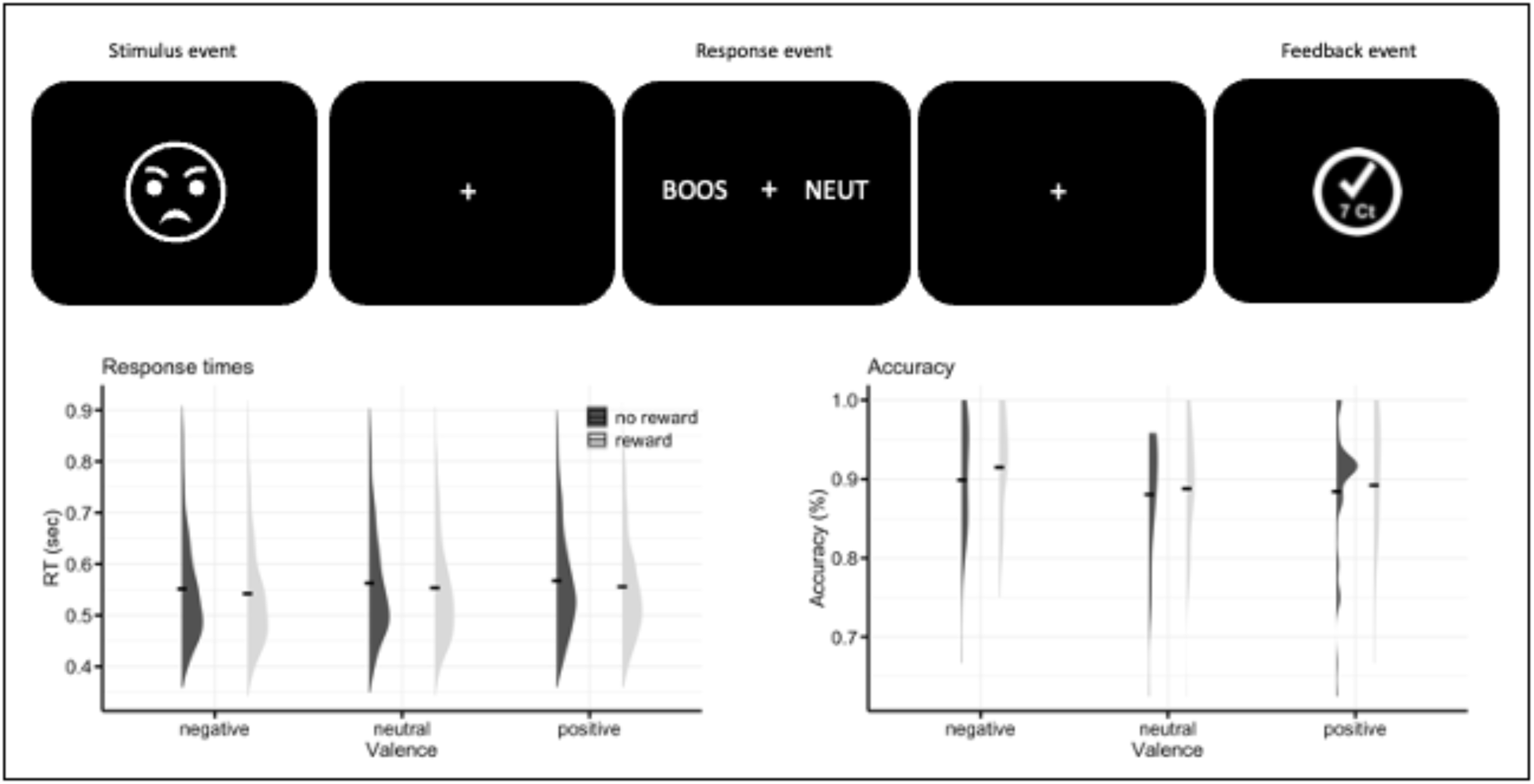
(A) Exemplary trial of the rewarded emotion-discrimination task paradigm, featuring a stimulus, a response, and a feedback event. Participants are asked to discriminate the emotional expression of the face by selecting the correct option during the response event. At the same time, the gender of the face signals whether a reward for a correct response in this trial. (B) Response times and accuracy according to the experimental factors reward (reward, no-reward) and Valence (negative, neutral, positive). The black line represents the mean values of RT and accuracy in the different conditions, the violin plots the distribution of the data. BOOS: Angry; NEUT: Neutral (Dutch). Note that the angry face emoticon is a <u>placeholder</u> and the real experiment used images of the NimStim face stimulus set (Tottenham et al., 2009), which includes photographs of actors expressing different emotions.

### 2.1.3 Image acquisition and pre-processing

MR images were acquired using a 3 Tesla Siemens Magnetom Prisma scanner (Siemens Medical Systems, Erlangen, Germany) with a 64-channel head coil at the Ghent Institute for Functional and Metabolic Imaging, Ghent University Hospital. For each functional run, we acquired T_2_*-weighted echo planar functional images in interleaved order with 69 transverse slices covering the whole brain, repetition time (TR) = 2090 ms, echo time (TE) = 27 ms, flip angle = 79°, field of view (FOV) = 112 mm, multiband acceleration factor = 3; no interslice gap and voxel size = 1.7 × 1.7 x 2 mm. T1-weighted structural volumes using a 3D magnetisation-prepared rapid acquired gradient echo (MP-RAGE) sequence (TR = 2250 ms; TE = 4.18 ms; FOV = 256 mm; flip angle = 9°; 176 sagittal slices; matrix size = 256 × 256; voxel size = 1 × 1 × 1 mm; duration = 5.14 min) were acquired for co-registration with functional scans. Additionally, a T2-weighted structural scan (TR = 3200 ms; TE = 4.08 ms; FOV = 230 mm; flip angle = 120°; 192 sagittal slices; matrix size = 256 × 256; voxel size = 0.4 × 0.4 × 0.9 mm; duration = 5.09 min) was acquired for the localization of the dopaminergic SN/VTA complex. Finally, a neuromelanin sensitive T1-weighted turbo spin echo (T1-TSE) scan (TR = 559 ms; TE = 98 ms; FOV = 192 mm; flip angle 1 = 70°; flip angle 2 = 180°; 10 sagittal slices; matrix size = 384 x 384; voxel size = 0.5 **×** 0. **×** 3 mm; duration = 9.51 min) was acquired for the localization of the LC. The total scanning time, which included a localiser scan, the MP-RAGE scan, a fieldmap, three functional runs, a T2 and a T1-TSE scan was approximately 60 min.

Statistical Parametric Mapping Software (SPM12, Wellcome Trust Centre for Neuroimaging, London, UK, http://www.fil.ion.ucl.ac.ak/spm/software) was used for image pre-processing. Functional scans were: (i) realigned and unwarped; (ii) slice time corrected to the middle slice in each repetition time; (iii) spatially normalized using the T_1_-weighted image to improve segmentation accuracy to fit the standardized MNI-152 template. Normalized images were resampled into 1.5 × 1.5 × 1.5 mm voxels for ROI analyses and 3 × 3 × 3 mm for the whole brain analysis (see Supplementary Materials), (iv) and smoothed using a 3.5 mm for ROI analyses and 8 mm for the whole brain analysis full width at half-maximum isotropic Gaussian kernel. FSL’s tool ICA-AROMA was used to correct for (MB exacerbated) motion artifacts (Kelly et al., 2012). Anatomical images (T1, T2, and T1-TSE) were co-registered to the mean functional image and normalized to fit the standardized MNI-152 template.

### 2.1.4 Behavioural analyses

All behavioural statistical analyses were performed in Rstudio (R version 4.2.2). Accuracy data (%) and response times (RT, correct responses only) averaged across all blocks were submitted to a repeated-measures analysis of variance (rANOVA) with the factors Reward (reward, no-reward) and Valence (negative, positive, neutral). Greenhouse-Geisser corrections are applied in case of sphericity violations. Significant interactions (and main effects of factors with more than two levels) were followed by post hoc t-test.

### 2.2 fMRI analysis

#### 2.2.1 First level model

Statistical parametric maps were calculated for each participant using a general linear model analysis within an event-related design. The high-pass filter was set to 128 seconds to remove low-frequency noise and each experimental condition was convolved with the canonical haemodynamic response function. Contrast images that represented the participant’s unique BOLD signal modulations in different conditions according to the experimental design (Reward x Valence) and trials events (stimulus, response, and feedback) were created and assigned to a total of 19 condition-specific regressors (including 1 regressor for incorrect trials). Finally, six motion regressors derived from the realignment procedure were included as additional regressors of no interest. Although the focus of this study is on modulations within LC and SN/VTA, we have included a whole-brain analysis to provide a for complete overview in the Supplementary Materials.

#### 2.2.2 ROI analysis

Individual masks for the LC and SN/VTA regions were manually drawn in MRIcroGL (Rorden & Brett, 2000) using the normalized single-subject T1-TSE and T2 scans respectively. The localization of LC and SN/VTA is illustrated on single-subject anatomical images in Figure 2A, along with the respective averaged mask across all participants. The LC and SN/VTA segmentation was guided by previous work (Düzel et al., 2009, 2015; Keren et al., 2009; Krebs et al., 2011, 2012, 2018). Note that we focused on the ventromedial part of the SN/VTA complex which has been considered particularly relevant for reward/valence processing (Düzel et al., 2015). The outer boundaries of the averaged bilateral masks in MNI coordinates and aligned to the brainstem axis are: LC (left/right x=-8/8; ventral/dorsal y=-36/-41; rostral/caudal; z=-17/-30); SN/VTA (left/right x=-13/14; ventral/dorsal y=-7/-20; rostral/caudal z=-8/-14). Mean parameter estimates (beta values) were extracted from the individual masks using the MarsBaR toolbox version 0.44 averaged across runs for each condition and converted to an SPM-compatible format. The beta values from each ROI were submitted to a rANOVA with three factors Reward (reward, no-reward), Valence (negative, positive, neutral), and Event (stimulus, response). Greenhouse-Geisser correction to degrees of freedom and *p* values was applied in case of sphericity violations. Significant interactions (and main effects of factors with more than two levels) were followed by post hoc t-test. The statistical analyses were performed in Statistical Package for the Social Sciences (SPSS), version 28.

**Figure 2.**
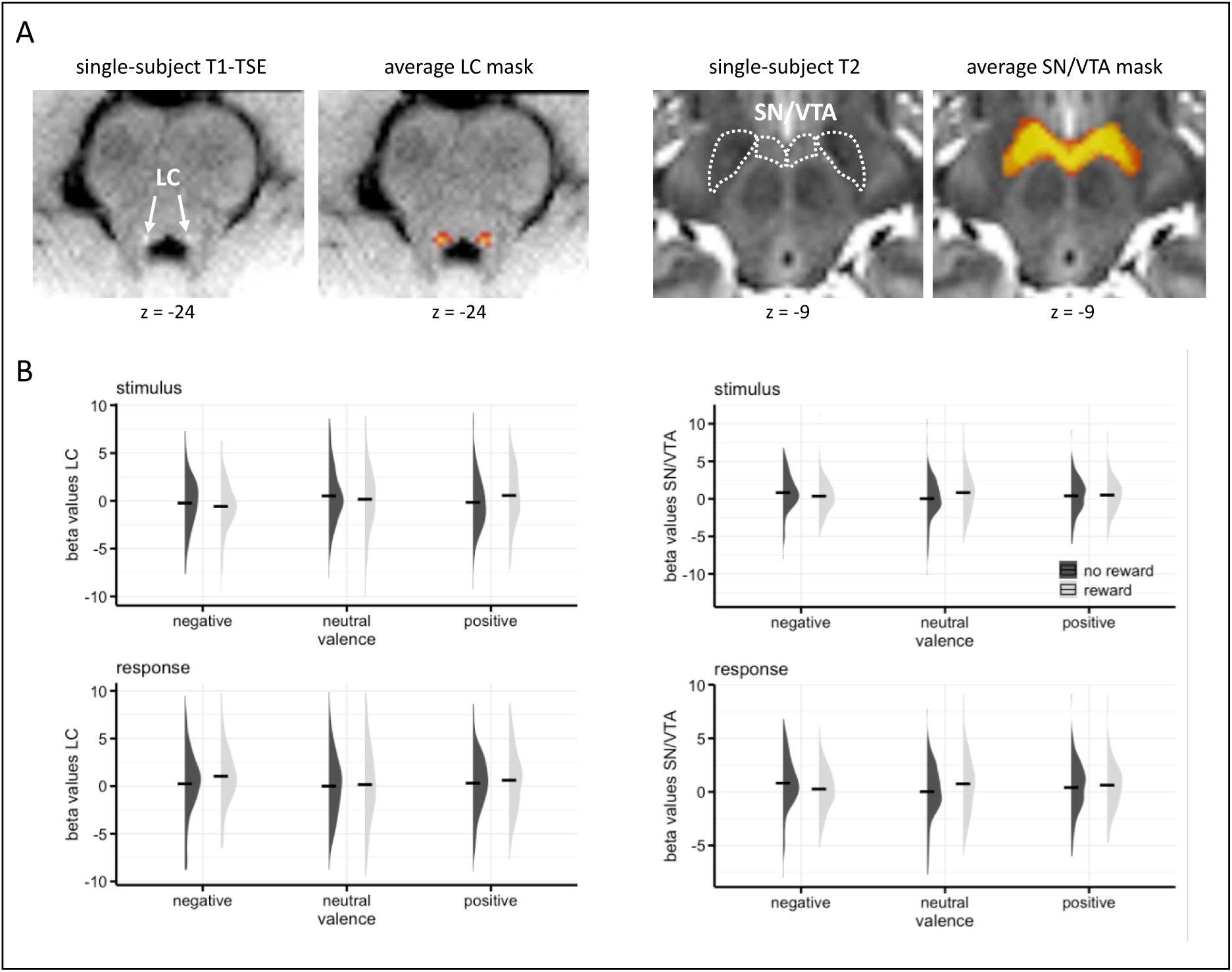
(A) An illustration of the LC (left) and SN/VTA (right) masks of an individual and group average. (B) Mean beta values for each condition in the stimulus (top) and response condition (bottom) of the LC (left) and SN/VTA (right). The black line represents the mean values of RT and accuracy in the different conditions, the violon plots the distribution of the data.

#### 2.2.3 Generalized Psychophysiological Interactions (gPPI) analysis)

To investigate the connectivity modulated in a task-dependent manner between predefined seed regions (LC and the SN/VTA) and other regions in the brain, we used a gPPI analysis (McLaren et al., 2012) implemented in CONN toolbox in Matlab (version 2022b). The gPPI analysis convolves the BOLD signal with the canonical hemodynamic response function for each condition before defining the contrast, creating a unique psychological regressor for each condition (unlike the standard PPI approach, which includes contrast information when forming a psychological regressor). To determine the interaction effects between the regressors (gPPI term, physiological vector, and psychological vector), random-effects analyses were performed for those contrasts that featured significant differential activity in the ROI analysis (*p* < 0.05, FDR corrected). In addition, covariations between LC and SN/VTA for both experimental factors were explored employing a direct ROI-to-ROI approach.

## 3. Results

### 3.1 Behavioural results

Mean accuracy rates for negative (M = 0.91, SD = 0.29, positive (M = 0.89, SD = 0.32) and neutral valence trials (M = 0.88, SD = 0.32) did not differ significantly (main effect Valence F(2,74) = 2.72, *p* = .066). Mean accuracies for reward (M = 0.90, SD = 0.31) and no-reward trials (M = 0.89, SD = 0.32) did not differ significantly either (main effect Reward F(1,37) = 1.15, *p* = .211), and there was no interaction between the factors (*p* > .9).

In the RT data, we did observe a main effect of Valence (F(2,74) = 7.80, *p* < .001), with significantly shorter RTs for negative (M = 546 ms, SD = 108) compared to both positive (M = 560 ms, SD = 108; t(71) = -3.94, *p* < .001 Bonferroni corrected) and neutral valence trials (M = 556 ms, SD = 109 t(71) = -2.75, *p* = .015 Bonferroni corrected). Further, there was a main effect of Reward (F(1,37)=10.34, *p* = .001), with shorter RTs for reward (M = 549 ms, SD = 107) compared to no-reward trials (M = 559 ms, SD = 110). There was no interaction between the factors (F(2,74) = 0.09, *p* = .917). Performance data are shown in Figure 1B.

### 3.2 fMRI results

#### 3.2.1 ROI analysis

An overview of the ROI analysis results can be found in Table 1. The rANOVA of beta values in the LC revealed a trend for the factor Reward (F(1,37) = 3.64, *p* = .064) with higher activity in reward compared to no-reward trials. This trend was accompanied by a significant interaction between Reward and Valence (F(2,74) = 3.79, *p* = .027). Post hoc tests showed that this interaction was due to significantly increased activity for positive reward trials compared to both neutral reward trials (t(37)= 2.68, *p* = .011) and positive no-reward trials (t(37)=3.32, p=.002). In addition, the analysis revealed a significant interaction between Valence and Event (F(2,74) = 5.11, *p* =.008), which arose from differentially increased activity for negative valence trials in the response phase in comparison with neutral (t(37) = 1.87, *p* = .007) and positive valence (t(37) = 1.35, *p* = .041). In contrast, during stimulus presentation, negative valence was associated with reduced activity compared to positive valence (t(37) = -2.75, *p* = .009). Consistent with this reversal, activity for negative valence was significantly lower in the stimulus compared to the response phase (t(37) = -3.13, *p* = .003). The main effects of Valence and Event were not significant (both *p* > .200) beyond the above interactions. The remaining two interactions were not significant (both *p* > .100).

**Table 1.**
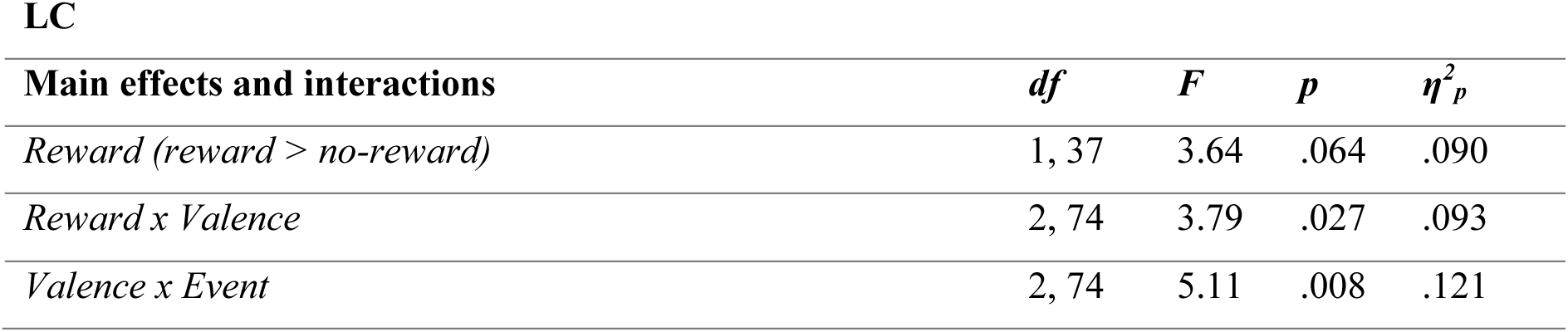
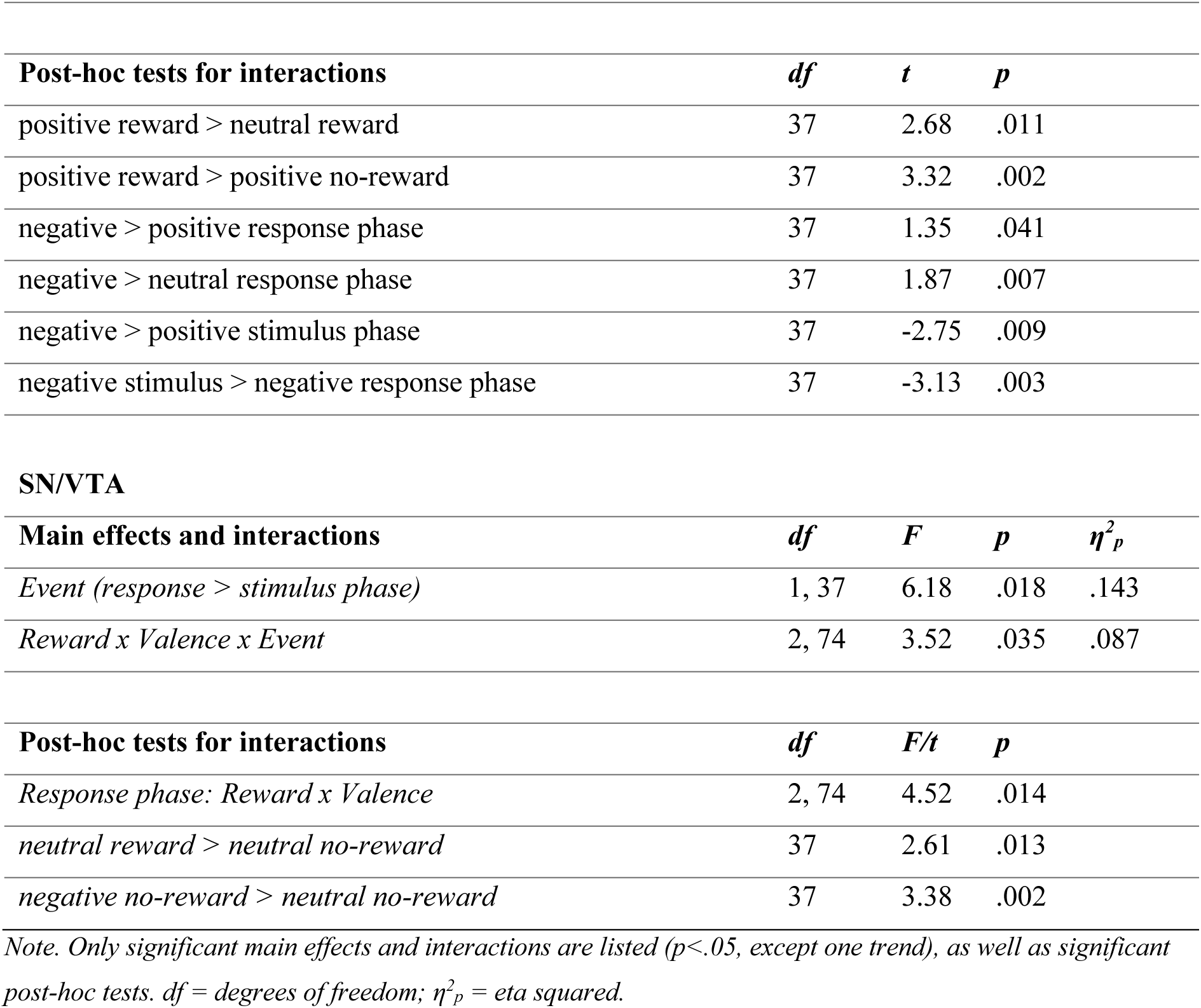
Significant effects in the ROI analysis in LC and SN/VTA.

The corresponding analysis in the SN/VTA revealed a significant main effect of Event (F(1,37) = 6.18, *p* = .018) with globally increased activity for the response compared to the stimulus event. This main effect was embedded in a significant three-way interaction between Reward, Valence, and Event (F(2,74) = 5.52, *p* = .035). Breaking down this interaction by Event type revealed a significant two-way interaction between Reward and Valence for the response event (F(2,74) = 4.52, *p* =.014), but not for the stimulus event (*p* > .200). The interaction in the response phase was characterized by increased activity for neutral reward compared to neutral no-reward trials (t(37) = 2.61, *p* = .013), as well as for negative no-reward compared to neutral no-reward trials (t(37) = 3.38, *p* = .002). The main effects of Reward and Valence were not significant (both *p* > .400) beyond the above three-way interaction. The three remaining interactions were not significant (all *p* > .100).

#### 3.2.2 gPPI results

We also explored condition-dependent covariation patterns between the SN/VTA and LC seed regions and the rest of the brain via gPPI analyses. Note that only comparisons that yielded significant effects in the above ROI analyses were considered. An overview of significant whole-brain covariations can be found in Table 2. For the contrast based on the Reward x Valence interaction, we found that across events, positive reward trials elicited connectivity between LC and right lateral occipital cortex (*p* < .025) compared to the neutral reward trials. No significant covariations were found between the LC and other regions for positive reward trials compared to positive no-reward trials (across event type). For the contrasts based on Valence x Event interaction, we found significant covariation between the LC and several clusters in the occipital cortex. Specifically, contrasting negative versus positive valence trials revealed negative covariations with the left and right occipital pole and right lateral occipital cortex in the response phase (left occipital pole: *p* = .006; right occipital pole: *p* < .001; right lateral occipital cortex: *p* < .001). The contrast negative versus neutral valence revealed a positive covariation with the right occipital pole (*p* < .001). No further significant covariations were observed for the LC or SN/VTA on the whole-brain level (all *p* > .05).

**Table 2.**
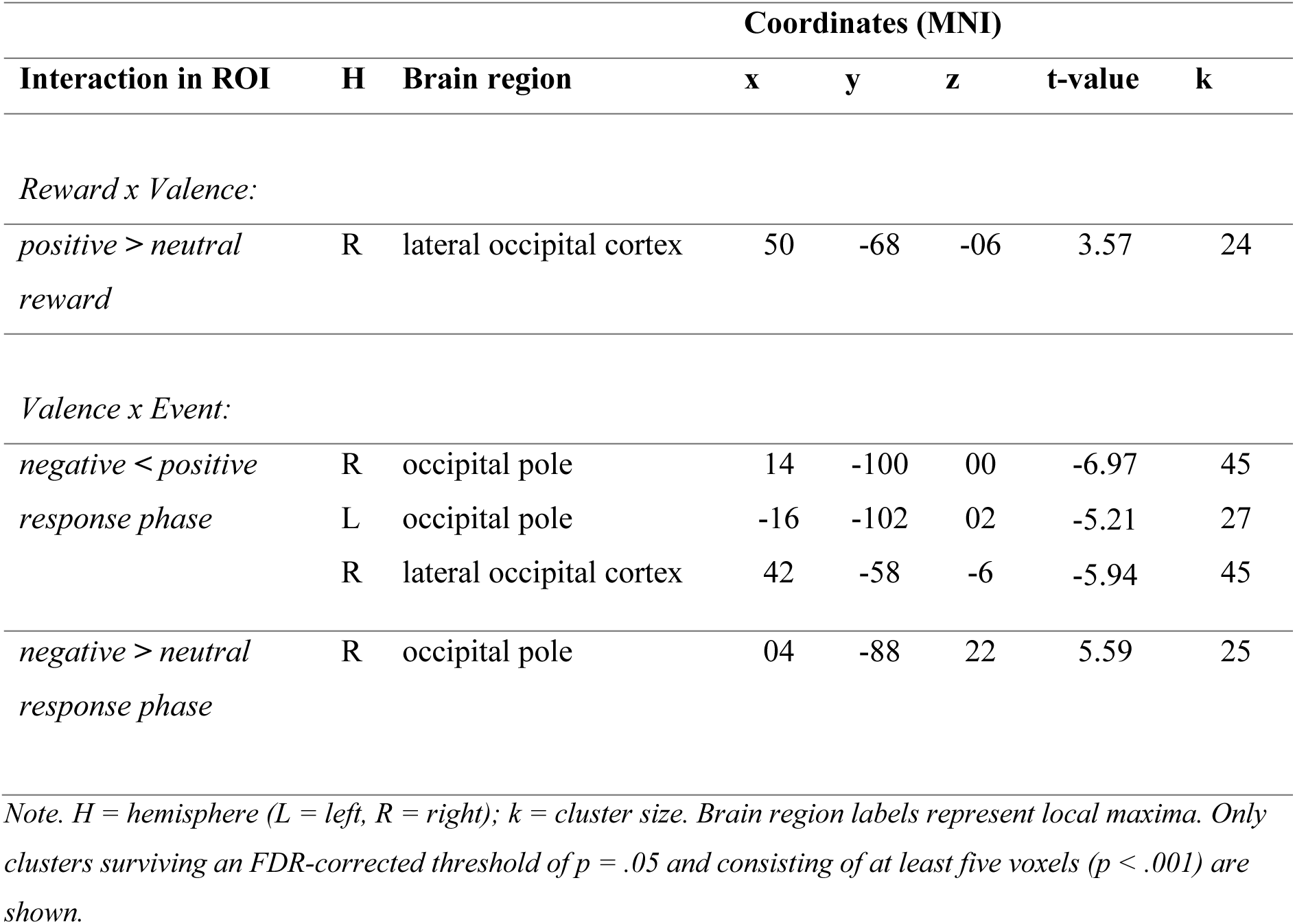
Significant generalized psychophysiological interactions (gPPI) between the LC and the whole brain.

Finally, we assessed the connectivity between LC and SN/VTA directly (ROI-to-ROI approach) to explore potential concerted modulations during reward and valence processing. This analysis revealed increased connectivity between LC and SN/VTA for reward compared to no-reward trials (t(37) = 2.36, *p* = .024) in the response phase. There was no covariation related to the stimulus condition, or to the valence contrasts (negative vs. neutral; negative vs. positive, positive vs. neutral) in either the stimulus or response condition.

## Discussion

While animal studies have shown overlapping responses of the DA and NA systems to behaviourally relevant stimuli, modulations in these systems in response to different stimulus types have predominantly been studied independently from each other in humans (but see this study on conflict processing (Köhler et al., 2016)). The present study aimed to investigate the distinct and joint contributions of the SN/VTA and LC activity during reward and emotional valence processing in one experimental design. This design also allowed us to explore potential interactions between reward and emotional valence. Behaviourally, we found that participants were significantly faster to respond to both negative valence and reward-predicting stimuli compared to their respective neutral and no-reward counterparts. This pattern replicates the well-known motivational effect of reward cues on the one hand (Braver et al., 2014), as well as the arousing effect of negative saliency on the other hand (Carver & Harmon-Jones, 2009). Moreover, this pattern indicates that response facilitation in this task is driven by the specific stimulus type (negative or reward-predicting) rather than general positive valence, which would be shared between reward-predicting and positive stimuli.

On the neural level, we found that LC activity was higher for trials entailing reward prospect (*p* = .06). This is in line with animal studies demonstrating that noradrenergic cells are sensitive to reward contingencies (Bouret and Richmond, 2010; Bouret et al., 2012; Bouret et al., 2015; Jahn et al., 2018). Importantly, this sensitivity to reward was qualified by emotional valence (reward x valence interaction), with a differential boost for reward-predicting positive valence trials as compared to reward prospect or positive valence alone. This could be interpreted as an additive effect of the two stimulus dimensions due to shared (or *congruent*) valence (Park et al., 2019). Notably, this effect (boost for reward-predicting positive trials) was observed across the trial, i.e., not distinct for either the stimulus or response phase. Finally, we also found a significant relative *increase* of LC activity for negative valence in the response phase, and a relative *decrease* for negative valence in the stimulus phase. The increase of activity in the response phase is in line with the LC’s sensitivity to aversive events in general (Aston-Jones & Cohen, 2005; Sara & Bouret, 2012), and also corresponds to the behavioural facilitation we observed. What is less intuitive, is the relative activity decrease for negative valence during stimulus encoding. One possible interpretation could be that the negative valence is not driving LC activity as long as no action is required. This would however be in contrast with the involvement of the noradrenergic system in response to encountering infrequent stimuli in an oddball paradigm (Nieuwenhuis et al., 2005). However, it is possible that in the current paradigm the interaction between different types of salient stimulus dimensions, as well as the separation of stimulus and response phase give rise to a more nuanced activation pattern in LC – which would also accommodate the increased activity for positive reward trials (or valence-congruency effect) in LC described above. An alternative interpretation could be related to response inhibition: since participants are instructed to not react until the response options are presented on the screen, LC activity, which may otherwise increase in the presence of negative valence, might be suppressed in the stimulus phase. One possible mechanism that may regulate these selective arousal states are the GABA neurons in the peri-LC. These neurons, which receive descending input from e.g., the mPFC (Lu et al., 2012), directly inhibit the LC, and recent work in mice has shown that in response to salient stimuli, the GABA neurons increase their activity (Luskin et al., 2022). Moreover, activating peri-LC GABA neurons decreases exploratory behaviour. In turn, increased activity for negative stimuli during the response phase might reflect resource allocation needed for responding to a behaviourally relevant event – similar to modulations observed in rewarded motor tasks (Bouret & Richmond, 2015). Since the movement itself (pressing the correct button) was the same across conditions, different salient outcomes (earn a reward, increase vigilance in the face of a potential threat) could increase the mobilisation energy regulated by NA.

Generally, investigations into the involvement of the LC in cognitive processes in humans are still scarce – which is partly due to challenges associated with scanning brain regions in the in the vicinity of ventricles and vessels (see below). The current results extend previous findings in other domains, including reward-based decision making (Varazzani et al., 2015), and the processing of infrequent (Krebs et al., 2018) or conflicting events (Hu et al., 2013; Köhler et al., 2016). What unifies these situations or stimuli is that they all are behaviourally relevant or at least salient with respect to other events in the respective task context. Moreover, the present LC modulations show that different sources of salient information can be integrated into an additive effect – in this case reward and positive valence.

Activity in the SN/VTA revealed differential modulations during the response phase only – i.e., when actions had to be selected and implemented. Specifically, activity in the response phase was differentially increased for reward compared to no-reward trials of neutral valence, but interestingly, also for negative compared to neutral valence trials that entailed no reward prospect. While the first of these observations (sensitivity to reward) replicates a well-established phenomenon in both human and animal neuroscience (e.g., Düzel et al., 2009; Schott et al., 2008), the latter (sensitivity to negative emotional events) has not gained much attention in human neuroimaging studies. However, it is consistent with a more global role of the DA system in processing behaviourally salient stimuli, including reward-predicting, aversive, as well as surprising events (Bromberg-Martin et al., 2010). The observation of two almost independent modulations in the present data mirror the performance effects and seem to be hinting at coding *relative saliency,* rather than an additive (or congruency) boost. While speculative, it could be possible that different SN/VTA neuron populations are coding for these different forms of saliency (see e.g., Krebs et al., 2009).

Finally, we explored functional covariations of LC and SN/VTA across the whole brain based on significant contrasts in the ROI analysis. Notably, the only covariations we found were between LC and occipital areas. Specifically, we found increased connectivity between LC and right occipital cortex during positive compared to neutral valence trials in the reward condition, independent of the phase (mirroring the prioritization of the positive reward condition observed in the LC ROI analysis). Moreover, connectivity was increased between LC and right occipital pole for negative compared to neutral valence in the response phase, independent of reward (mirroring the prioritization of negative valence especially during the response phase). These occipital regions are involved in visual processing general, and in particular subserving object processing (including faces) and visuo-spatial processing (Haxby et al., 1991). The increased connectivity between LC and lateral occipital cortex for positive reward-predicting as well as negative faces aligns with the finding that pharmacological up-versus down-regulation of noradrenergic signalling in humans improved versus impaired attentional selection and discrimination in lateral occipital cortex (Gelbard-Sagiv et al., 2018). In addition, we found a *negative* covariation between LC and bilateral occipital pole and right lateral occipital cortex when contrasting negative to positive valence in the response phase (again independent of reward). We tentatively suggest that this could be related to dampening of connectivity between LC and occipital areas due to higher cognitive load imposed by negative emotions, leading to network reconfiguration (Fraser et al., 2012), or an attentional bias where resources are reallocated to focus on internal emotional states or response preparation. note that these analyses were exploratory and partly contradictory and hence call for replication. Finally, the ROI-to-ROI connectivity analysis revealed positive covariation between LC and SN/VTA for reward compared to no reward trials in the response phase. This resonates with their reciprocal neuronal connections (Chandler et al., 2014), which may give rise to the invigoration of actions to that hold rewarding outcomes (Bouret & Richmond, 2015). Regarding the directionality of the influence, on the one hand it has been suggested based on animal models that the VTA triggers phasic activation of the LC in response to reward-predicting stimuli, which in turn serves further allocation of attention (Hofmeister & Sterpenich, 2015). On the other hand, animal addiction models posit that LC inputs to the VTA can modulate drug-seeking behaviour (Solecki et al., 2022) – which relies on the same dopaminergic circuits as reward processing. While these are interesting avenues for further research, the present data is considered merely exploratory in this regard.

Previous work on the role of the NA and DA source regions in processing reward-predicating and/or valenced stimuli in humans have mostly been selective for one of the systems or one of the stimulus dimensions. In the present study we observed contributions from both regions to reward and valence processing within the same paradigm. First and foremost, we saw that especially negative valence stimuli were associated with increased activity in both LC and SN/VTA during the response phase. This confirms well-known sensitivity of the LC for aversive stimuli (Sara & Bouret, 2012). In contrast, sensitivity of the SN/VTA for negative stimuli is less well established but has been put forward in animal models (Bromberg-Martin et al., 2010; Horvitz, 2000). As such, the present data confirm that both systems are especially sensitive to behaviourally relevant *negative* valence stimuli – as one type of salient stimuli in the environment. Considering the other salient stimulus dimension, i.e., reward prospect, the patterns were slightly diverging in the two regions: while the LC featured a valence-dependent boost in reward trials with highest activity for positive valence, the SN/VTA featured selective boosts for reward trials of neutral valence as well as for negative valence trials without reward prospect. The LC pattern is consistent with the notion of an additive (or congruency) effect of overlapping valence. Specifically, in a previous study we found that signalling reward prospect by positive valence explicitly led to behavioural facilitation (Park et al., 2019). In the present study, however, there was no additional performance facilitation, which is likely due to the separation of stimulus encoding and response implementation. In contrast to LC, the SN/VTA appears to code trials in terms of relative saliency (reward or negative valence) but does not “perform” an integration at this stage in the present paradigm. It is likely that such integration is realized in higher-order regions, including the striatum and fronto-parietal cortex, as we have seen in our previous study (Park et al., 2019). Together, the prioritization of reward and negative valence stimuli independently in the SN/VTA - together with the distinct LC modulation for negative valence stimuli - are most consistent with the two independent main effects observed in task performance. Although the reward-related modulations the present study are generally consistent with related animal work that looked at both dopaminergic and noradrenergic cells within the same task (Bouret et al., 2015), a direct comparison seems challenging as we added an additional valence dimension here.

Animal work has shown that DA and NA activity differs: DA responses are stronger when a reward is anticipated, whereas NA responses increase for unrewarded actions (Bouret et al., 2012). An interesting general observation of the current study is that the modulations in both regions appear to be better aligned to the response phase following reward prospect rather than to the presentation of the reward-predicting stimulus itself. The same holds for modulations in response to negative valence. Importantly, the increase in activity in the response phase is not a mere reflection of motor implementation as it is selective for specific conditions. Moreover, the response phase entails more than implementing a response: participants select the appropriate response from two (conflicting) valence options, hence involving a more or less effortful decision process. In contrast, the stimulus phase mainly required participants to the encode the two stimulus dimensions (reward and valence). Thus, the concerted activation of the DA and NA system may be particularly important to select and invigorate responses to behaviourally relevant stimuli rather than mere stimulus evaluation (Hofmans et al., 2022). As such, this novel paradigmatic distinction between stimulus encoding and response implementation sheds new light on the contributions of the LC and SN/VTA during the processing of reward prospect and valence.

Thus, our data support the notion that both LC and SN/VTA are involved in reward and emotional valence processing. Both regions seem to be mostly sensitive to reward-predicting as well as negative events. However, differences between regions surfaced based on the trials phase, as well as when considering interactions between the different factors. While the LC featured an additional boost of the reward response by positive emotional valence, activity in SN/VTA was independently modulated by reward and emotional valence, aligning more with the coding of relative saliency. Hence, despite the overlap, they do likely fulfil slightly different roles, which might be related to their respective projection targets (Ranjbar-Slamloo & Fazlali, 2019). Previous work has shown saliency engages other regions such as the anterior insula and anterior cingulate cortex, ventromedial prefrontal cortex, amygdala, ventral striatum, thalamus, and occipito-temporal cortex. While our whole-brain results (see Supplementary data) did show involvement of cortical structures such as the occipital cortex and medial frontal cortex, only the occipital cortex covaried with LC activity in the connectivity analysis (see above).

In this final section we are considering limitations of the present study. First, neuroimaging of small, deep structures requires smaller voxel-sizes for accurate localization and signal extraction, which in turn comes with the inherent limitation of a low signal-to-noise ratio (Edelstein et al., 1986). Hence, we might be underestimating, or even missing, potential existing modulations in the target regions. Moreover, the increased MB sampling frequency may induce noise amplification, especially in subcortical regions (Risk et al., 2021), but deemed necessary to acquire the ambitious sequence scanning both the brainstem with high resolution as well as the neocortex to allow for connectivity analyses. Another challenge is the close vicinity of LC and SN/VTA to adjacent ventricles and vessels, rendering them susceptible to physiological noise. To account for these interferences, we have applied ICA-AROMA to the functional data, which is a data-driven method to identify and remove motion artefacts stemming from head movements but also blood pulsation. Due to a technical artifact most likely induced by the interaction between the MB sequence and (head) motion, we needed to exclude six participants, and therefore did not reach our pre-registered sample. Further, it was surprising that no significant covariation clusters were detected when using SN/VTA as a seed region. This could be related to the signal-to-noise issue but might be further complicated by functional segregation within this region. Specifically, it has been suggested that different neuron populations in the SN/VTA complex subserve different functions (Düzel et al., 2009), which could dilute results from (potentially underpowered) connectivity analyses. That said, further segregation of the SN/VTA is beyond the scope of the present study and would require a larger sample. Finally, in this article, we have considered the activity of LC as a proxy for NA. However, these neurotransmitters are synthesized from the same precursor, tyrosine, and undergo similar biochemical transformations, and the LC also co-releases dopamine (Kempadoo et al., 2016). Hence, the interaction between NA and DA in the LC may also have played a crucial role in modulating the studied processes. The coexistence and interaction of these neurotransmitters suggest a complex neurochemical balance within the LC, potentially contributing to its multifunctional role in brain regulation. Further studies are needed to elucidate the precise mechanisms and implications of dopaminergic and noradrenergic interactions in this and other brain structures. The present results should therefore be replicated and confirmed in future studies.

To conclude, the current study explored the intricate roles of the LC and SN/VTA system in the processing of reward and emotional valence. The findings underscore the sensitivity of both LC and SN/VTA to reward prospects as well as to negative valence. These findings challenge the dominant view of the SN/VTA involvement in merely positive events. Moreover, we found evidence for an *additive effect* (boost) of positive valence and reward in the LC, while responses in the SN/VTA were more aligned with coding *relative saliency* of reward and emotional events in their respective context. Intriguingly, both regions exhibited strongest modulations during the response phase, thereby emphasizing their essential role in goal-directed action invigoration, above and beyond stimulus encoding. However, the replication and validation of these findings in future studies are imperative to fortify our understanding of the interplay of DA and NA underlying reward and emotional processing.

