## Supplementary Materials for "Exploring distinct and joint contributions of the Locus Coeruleus and the Substantia Nigra/Ventral Tegmental Area complex to reward and valence processing using high-resolution fMRI"

**Whole brain analysis**

Voxel-wise random-effects analyses were performed for the reward emotional recognition paradigm using a 3 × 2 flexible factorial design, with Valence (positive/neutral/negative) and Reward (reward/no reward) as factors. For the Valence factor, we compared neutral to both positive and negative trials, as well as the differences between positive and negative trials in both directions. Activity maps for Event Type (stimulus and response phase) were submitted separately to the second level analysis. We also investigated potential interactions between the factors.

Activations were considered significant if they survived a cluster-level family-wise error (FWE) correction of *p* <. 05, based on an uncorrected voxel-wise threshold of *p* < .001.

**Results**
Significant activations were found for the response phase for the reward and valence condition (see Table SX), but no interactions between the factors was observed.

In the stimulus phase, no significant activations were found for the main effects of Reward and Valence, nor any interaction effects (all *p* > .05 FWE corrected).

**Table S1 Whole brain results in the response phase**

| Main effect |  |  | Coordinates (MNI) | | |  |  |
| --- | --- | --- | --- | --- | --- | --- | --- |
|  | **H** | **Brain region** | **x** | **y** | **z** | **t-value** | **k** |
| Reward |  |  |  |  |  |  |  |
| *Reward vs. No Reward* | R | Insula | 30 | 23 | -1 | 5.55 | 88 |
|  | R | Precuneus | 15 | -64 | 47 | 4.47 | 257 |
|  | R | Mid Cingulum | 6 | 20 | 41 | 3.88 | 88 |
|  |  | Pons | 0 | -31 | -34 | 5.20^#^ | 54 |
|  | L | Insula | -45 | 11 | -1 | 3.65^#^ | 54 |
| Valence |  |  |  |  |  |  |  |
| *Positive vs. Neutral* | R | Supramarginal Gyrus | 63 | -28 | 29 | 4.91 | 169 |
|  | L | Mid Temporal Gyrus | -39 | -61 | 14 | 4.73 | 65 |
|  | L | Anterior Cingulum | -3 | 35 | 8 | 4.51 | 148 |
|  | L | Mid Cingulum | -6 | -34 | 47 | 3.86 | 58 |
|  | R | Mid Cingulum | 12 | -28 | 38 | 4.48^#^ | 56 |
| *Negative vs. Neutral* | L | Supramarginal Gyrus | -63 | -49 | 29 | 4.97 | 109 |
|  | L | Anterior Cingulum | -6 | 35 | 5 | 4.21 | 61 |

*Note:* H: hemisphere; R = right; L = left

^#^ *p* < .01 (pons: *p* = .072; Insula: *p* = .072; Mid Cingulum = *p* = .064)
